# HiCDiff: single-cell Hi-C data denoising with diffusion models

**DOI:** 10.1101/2023.12.01.569684

**Authors:** Yanli Wang, Jianlin Cheng

## Abstract

The genome-wide single-cell chromosome conformation capture technique, i.e., single-cell Hi-C (ScHi-C), was recently developed to interrogate the conformation of the genome of individual cells. However, single-cell Hi-C data are much sparser and noisier than bulk Hi-C data of a population of cells, making it difficult to apply and analyze them in biological research. Here, we developed the first generative diffusion models (HiCDiff) to denoise single-cell Hi-C data in the form of chromosomal contact matrices. HiCDiff uses a deep residual network to remove the noise in the reverse process of diffusion and can be trained in both unsupervised and supervised learning modes. Benchmarked on several single-cell Hi-C test datasets, the diffusion models substantially remove the noise in single-cell Hi-C data. The unsupervised HiCDiff outperforms most supervised non-diffusion deep learning methods and achieves the performance comparable to the state-of-the-art supervised deep learning method in terms of multiple metrics, demonstrating that diffusion models are a useful approach to denoising single-cell Hi-C data. Moreover, its good performance holds on denoising bulk Hi-C data.

## 1. Introduction

The information about the three-dimensional (3D) conformation (structure) of a genome is important for analyzing and understanding its function such as gene regulation, enhancer-promoter interaction, and genome methylation. Hi-C is a widely used high-throughput next-generation sequencing assay for measuring pair-wise contacts between any pair of genomic loci (Pal, et al., 2019). Hi-C chromosomal contact data have revealed important genome conformation features such as topologically associated domains (TADs) and chromatin compartments in both a population of (bulk) cells (Dixon, et al., 2012; Lieberman-Aiden, et al., 2009; Nora, et al., 2012) and single cells (Nagano, et al., 2013; Nagano, et al., 2017; Tan, et al., 2018).

However, Hi-C data usually contain substantial noise due to multiple factors. The most common one is that the amplification step for the library preparation in the experiments introduces the distance-dependent amplified bias such that a higher noise-to-signal ratio against genomic distance exists. The restriction enzymes of cutting contacted DNA fragments off from the genome also have a biased preference, leading to over- or under-representation of some contacts. It is important to remove the noise in Hi-C data, particularly in very sparse single-cell Hi-C data with very low signal-to-noise ratio, in order to use them well in the downstream applications (Li, et al., 2015; Lieberman-Aiden, et al., 2009; Servant, et al., 2015; Wolff, et al., 2022).

Hi-C paired-end reads are usually mapped to a reference genome and then converted into chromosomal contact matrices/maps (Lieberman-Aiden, et al., 2009; Trieu and Cheng, 2016). In a contact matrix (M), each entry *M*[*i, j*] contains the number of reads indicating the frequency of two fragments i and j being in contact. Several deep learning methods have been developed to impute and/or denoise Hi-C contact matrices, including HiCSR (Dimmick, 2020), DeepHiC (Hong, et al., 2020), HiCPlus (Zhang, et al., 2018), VEHiCLE (Highsmith and Cheng, 2021), and HiCARN (Hicks and Oluwadare, 2022) for processing bulk Hi-C data of a population of cells as well as SCHiCEDRN (Wang, et al., 2023) for processing both single-cell and bulk Hi-C data and Higashi (Zhang, et al., 2022) and DeepLoop (Zhang, et al., 2022) for processing single-cell Hi-C data. All these methods are supervised learning methods, which require both labeled data (e.g., cleaner data or noiseless data) and noisy input data to train them. However, the pairs of labeled and noisy data may not always be available. The method trained on one kind of noisy and labeled data may not be applicable to another kind.

Inspired by both the Denoising Diffusion Probabilistic Models (DDPM) (Ho, et al., 2020; Sohl-Dickstein, et al., 2015) and Denoising Diffusion Restoration Models (DDRM) (Kawar, et al., 2022) that have achieved success in denoising and generating images (Goodfellow, et al., 2014; Guo, et al., 2023; Karras, et al., 2017; Lehtinen, et al., 2018; Zhang, et al., 2017), we developed a diffusion model – HiCDiff that can work in both supervised and unsupervised modes to denoise Hi-C data of both single cells and bulk cells. In both modes, HiCDiff uses a parameterized Markov chain model trained to learn the transition from noisy data to cleaner data to reverse a noise forward diffusion process of gradually adding Gaussian noise to Hi-C data.

The conventional diffusion models including DDPM, DDRM or SR3 (Saharia, et al., 2022) used to denoise data in other domains all employed U-Net architectures with small modifications to parameterize the Markov chain model in the reverse diffusion process. In this work, we adapted a DDPM with U-Net as the baseline method for denoising Hi-C data. Moreover, we designed HiCDiff that employs a residual network architecture with DDPM to denoise the Hi-C data of either a single cell or bulk cells. HiCDiff consistently performs better than DDPM on several single-cell and bulk Hi-C test datasets in terms of multiple evaluation metrics in both the unsupervised and supervised modes. Compared to the non-diffusion deep learning methods, HiCDiff outperforms most of them and is comparable to the state-of-the-art method.

## 2. Methods

### 2.1 Denoising Diffusion Framework and Training Algorithms of HiCDiff

HiCDiff uses the framework of the Denoising Diffusion Probabilistic Models (DDPM) (Ho, et al., 2020) to model how noise is added into data and it can be removed. It is inspired by nonequilibrium thermodynamics and uses a Markov chain to model how uncorrupted data (*y*_0_) gradually diffuse into noise (*y*_*T*_) from time 0 to T in the forward process (***q***): *y*_0_ → *y*_1_ → … → *y*_*T*−1_ → *y*_*T*_ and how the noise (*y*_*T*_) is gradually transformed back to the true/target data (*y*_0_) in the inverse process (***p***): *y*_*T*_ → *y*_*T*−1_ → … → *y*_2_ → *y*_0_.

In the forward process, Gaussian noise is added to each time *t* − 1 to generate corrupted data at time *t* according to the distribution: 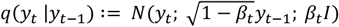, where is *β*_*t*_ is a variance schedule parameter at time *t* (1≤ *t* ≤ *T*) to control how much Gaussian noise is added at each time step *t*. Because the Markov property of the forward process, *y*_*t*_ can be directly sampled from *y*_0_ according to the distribution: 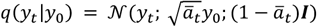 where 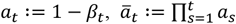. The forward process can be carried out straightforwardly to generate corrupted data to train a model to reverse the data corruption process iteratively.

*t*=0

In the reverse (denoising) Markov chain process: *y*_*T*_ → *y*_*T*−1_ → … → *y*_2_ → *y*_0_, the joint distribution of the data points: *p*_*θ*_(*y*_0:*T*_) can be calculated by the formula: 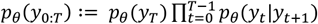, where *p*_*θ*_(*y*_*T*_) = *N*(*y*_*T*_; 0; *I*) and *p*_*θ*_(*y*_*t*_|*y*_*t*+1_) = *N*(*y*_*t*_; *μ*_*θ*_(*y*_*t*+1_, *t* + 1); *σ*_*θ*_(*y*_*t*+1_, *t* + 1)). *μ*_*θ*_ and *σ*_*θ*_ are the mean and variance of *y*_*t*_ that depends on *y*_*t*+1_. The mean can be predicted from *y*_*t*+1_ by a neural network-based generative model with parameter θ trained on the data generated in the forward process. In practice, because the mean is proportional to the difference between *y*_*t*+1_ and the added noise and *y*_*t*+1_ has been provided as input, a neural network (*f*_*θ*_) is usually trained to predict the Gaussian noise from *y*_*t*+1_ instead. Once the noise is predicted, the mean can then be calculated. *σ*_*θ*_ can be set to the same as the variance of time step *t* in the forward process. Then *y*_*t*_ can be sampled from the Gaussian distribution with the mean and the variance.

#### Algorithm 1

generate a noisy chromosomal contact matrix at time *t* in the forward process.

**Input**: *y*_0_, the noiseless chromosome matrix at the start time *t*_0_; *t*, a time step in the range [1, T]; *β*_*t*_, the noise level at time t based on a noise variance schedule in the range [0, 1]

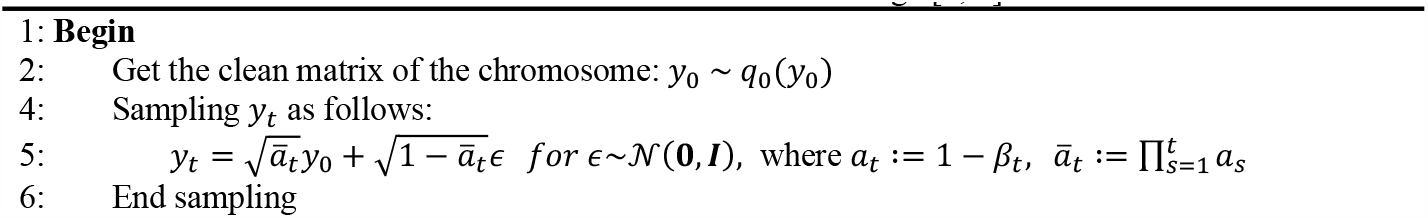
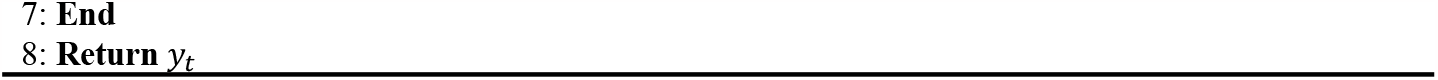

To apply the probabilistic diffusion framework above to Hi-C chromosomal contact matrices, **Algorithm 1** is used by HiCDiff to add Gaussian noise into a clean contact matrix (*y*_0_) to generate a corrupted matrix (*y*_*t*_) for any time *t* in the forward process. The corrupted contact matrices with the added noise are used to train a generative neural network to remove the noise in the reverse process.

The generative neural network model can be trained in either unsupervised learning mode (unconditional diffusion) or supervised learning mode (conditional diffusion). In the unsupervised model (Ongie, et al., 2020), the data (*y*_0_) is fully corrupted into noise (*y*_*T*_) in the *forward* process (**Figure 1A**), and then a generative model is trained to predict the noise to be removed from *y*_*t*_ to estimate the mean of *y*_*t*−1_. Once the model is trained, it can be used in the *inverse* process to recover *y*_0_ step by step, starting from the complete Gaussian noise *y*_*T*_ (Bardsley, 2012). The unsupervised mode is used when only presumably clean data (*y*_0_) is provided for training. **Algorithm 2** describes how a generative model (*f*_*θ*_) is trained in the unsupervised mode to predict the noise at any time step until it converges. In contrast, the supervised learning mode can be used when pairs of noisy data x and clean data *y*_0_ (target label) are provided to train the generative model. In this mode (**Figure 1B**), a noisy data matrix x (i.e., a condition) in conjunction with the corrupted data (yt) is used as input for a generative model to predict the noise to be removed to obtain the mean of *y*_*t*−1_ in the inverse process. **Algorithm 3** describes how a generative model (*f*_*θ*_) is trained in the supervised mode to predict the noise to be removed.

**Fig. 1.**
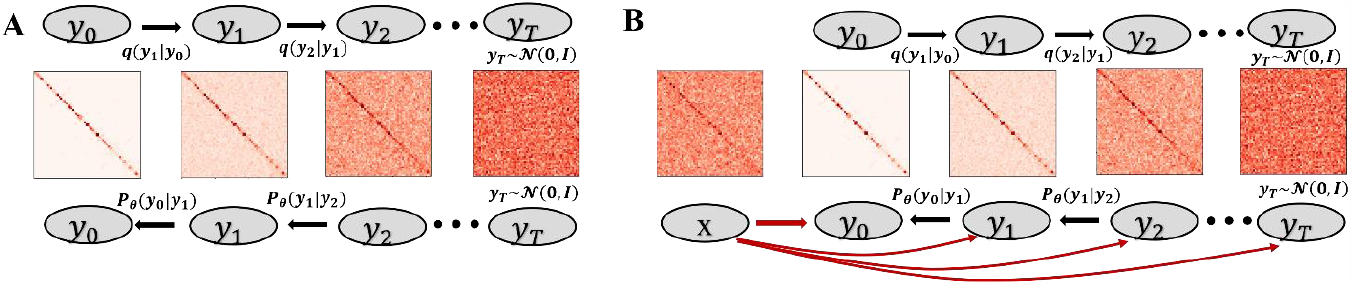
The forward and inverse processes of the HiCDiff for unconditional and conditional diffusion. **(A)** The unconditional diffusion model (unsupervised mode). In the forward process (*q*) at the top, the clean data (y0) is gradually corrupted until it becomes the complete noise (yT), and in the reverse process (*p*) at the bottom the noise is gradually removed until the clean data is recovered. The generative neural network model for removing noise is trained in the unsupervised mode. **(B)** The conditional diffusion model (supervised mode). The forward and inverse processes of the conditional diffusion are the same as those of the unconditional diffusion except that a conditional x, which is provided as an extra input, is concatenated with *y*_*t*_ at every time step t (red arrows) to train the generative neural network model to predict noise to be removed.

#### Algorithm 2

training a generative neural network model (*f*_*θ*_) in the unsupervised mode

**Input**: *y*_0_, the noiseless chromosomal matrix at the start time *t*_0_

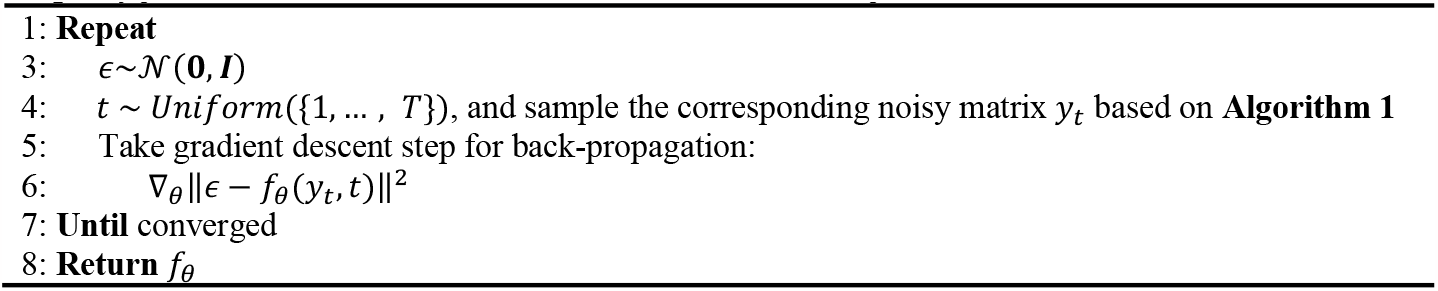

#### Algorithm 3

training a generative neural network model in the supervised mode

**Input**: *y*_0_, the noiseless chromosomal contact matrix at the start time *t*_0_; x, a noisy chromosomal contact matrix provided as a condition

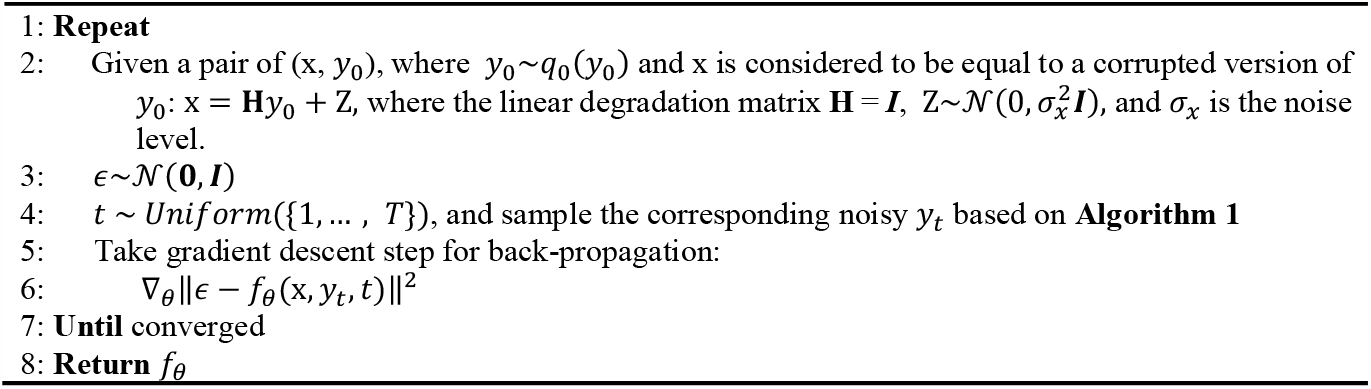

### 2.2 Inference algorithms for denoising chromosomal contact matrices in HiCDiff

The inference of HiCDiff uses the trained generative model in a reverse Markov chain process starting with Gaussian noise *y*_*T*_∼*N*(0, *I*) to generate cleaner data iteratively through *T* steps until recovering *y*_0_ according to the formula 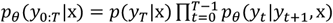, where *p*(*y*_*T*_|x): = *N*(0, *I*) and 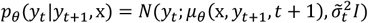. Here *μ*_*θ*_ is calculated from the prediction of the model (*f*_*θ*_) trained in either the supervised or unsupervised mode and 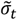 is calculated as 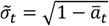, where 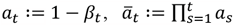 and *β*_*t*_ is a variance schedule parameter at time *t* used in the forward process **Algorithm 4** describes the process of denoising a contact matrix in the unsupervised mode based on the diffusion relationship between the original and spectral spaces, and **Algorithm 5** describes how to denoise a contact matrix in the supervised mode.

#### Algorithm 4

denoising a chromosomal contact matrix (x) in the unsupervised mode in the inference phase. ***Notations***: all the variables without bar (e.g., *y*_*t*_) denote the data (e.g., a chromosomal contact matrix) in the original space, and all the variables with bar (e.g., 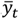) denote the data in the spectral space (the space of eigenvalues associated with a linear transformation matrix **H**). The variables with subscript *θ* connect the diffusion process between the original and spectral spaces. And a line started with **#** is a comment describing the following operations.

**Input:** a noisy chromosome contact matrix x, where the noise is controlled by the input noise level *σ*_*x*_; *f*_*θ*_, a trained model

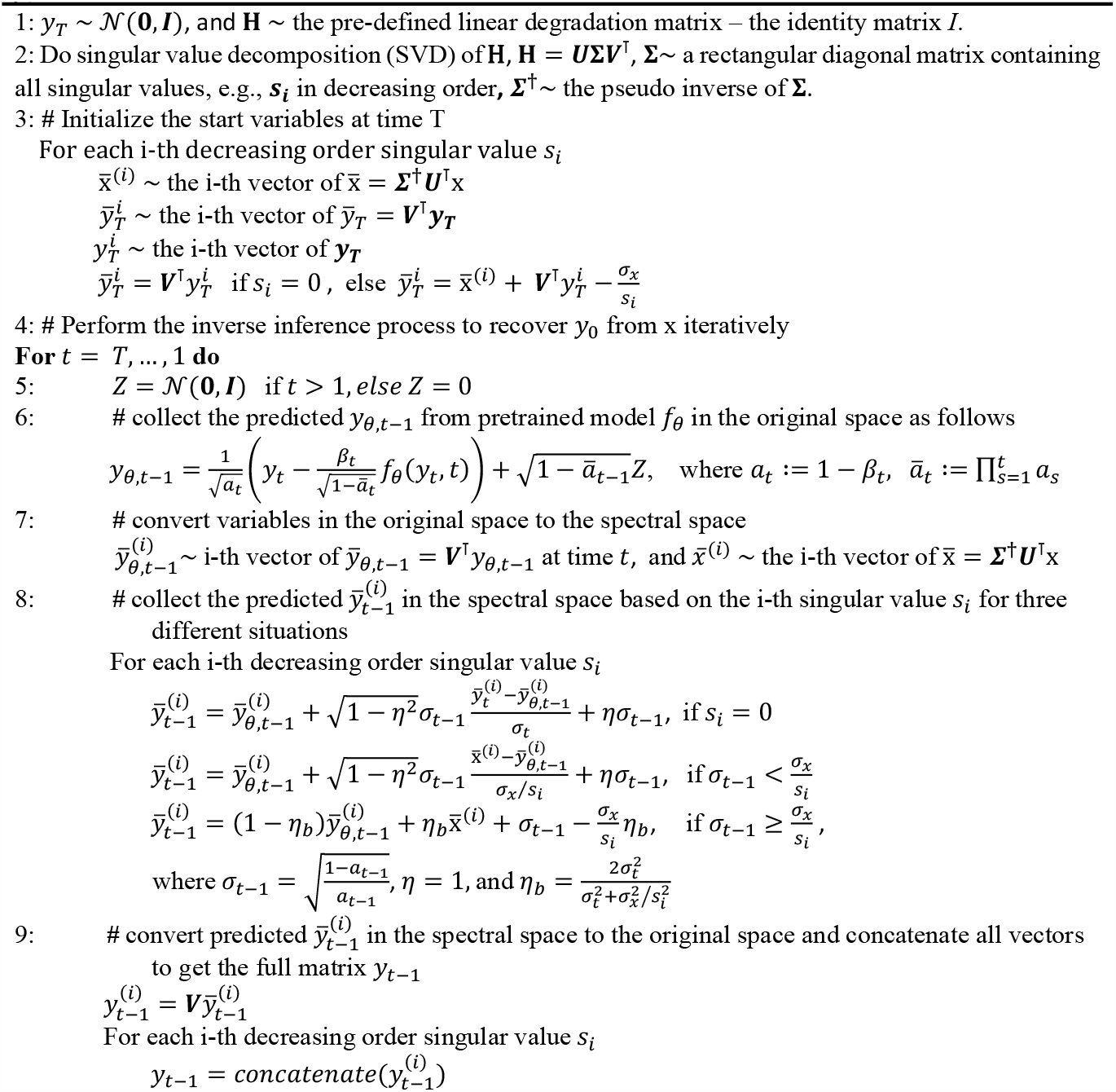
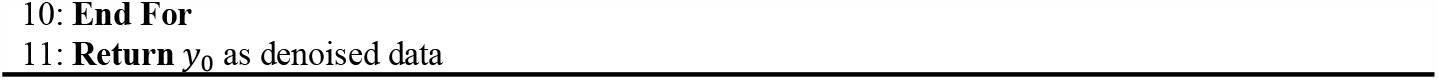

#### Algorithm 5

denoising a chromosomal contact matrix (x) in the supervised mode in the inference phase.

**Input**: x, a noisy chromosomal matrix; *f*_*θ*_, a trained model.

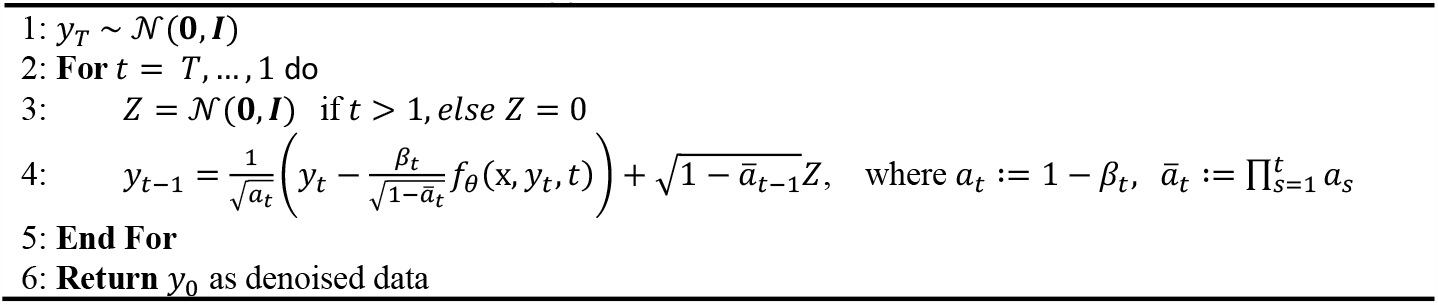

### 2.3 The deep learning architectures of generating cleaner data from noisier data in HiCDiff

We tested two different deep learning architectures to predict the noise that needs to be removed from the noisy data (*y*_*t*_) to obtain the mean of cleaner data (*y*_*t*−1_) with or without an input condition (x). One is the standard U-Net with multi-head attention layers used in the classic DDPM models (Ho, et al., 2020). This method is the baseline diffusion model, which is referred to as DDPM when compared with the other methods. Another one is a residual network illustrated in **Fig. 2**, which performs better than the U-Net and is used as the final generative model in HiCDiff. The residual network consists of 32 customized residual blocks with one 3×3 convolution layer preceding them and another 3×3 convolution layer following them, which is the same as the generator component used in a generative adversarial network method – SCHiCEDRN (Wang, et al., 2023) for denoising Hi-C data.

**Fig. 2.**
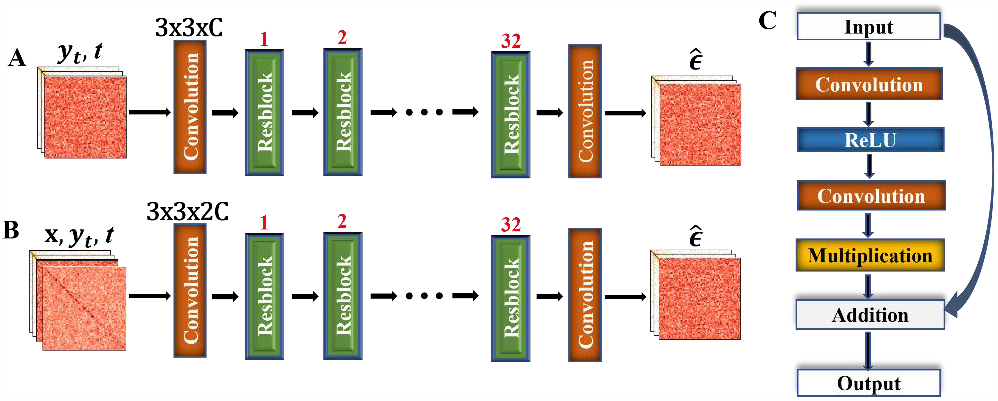
The architecture of the deep residual network used by HiCDiff to predict the noise 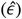 to be removed from a corrupted input matrix (*y*_*t*_) at time *t*. **(A)** The architecture of the unsupervised learning mode. It has one convolutional layer (filter size: 3 × 3 × C; C: the number of input channels), followed by 32 residual blocks (Resblock) and a final convolutional layer to predict the noise. **(B)** The architecture of the supervised learning mode. x is the conditional source matrix provided by users. (**C**) The layers of the residual block (Resblock).

Both the unsupervised (**Fig. 2A**) and supervised (**Fig. 2B**) HiCDiff use the same residual network architecture during the training and inference, except that the number of input channels of the first convolution layer of the supervised HiCDiff is twice that of the unsupervised HiCDiff because the former takes an extra condition matrix x as input in addition to the noisy data matrix *y*_*t*_ used by the latter. Once the unsupervised and supervised models (*f*_*θ*_) are trained by **Algorithm 2** and **3** respectively, they can be used by **Algorithms 4** and **5** respectively to generate the cleaner data (*y*_*t*−1_) iteratively in the reverse process of diffusion.

### 2.4 Datasets for training, validating, and testing HiCDiff

Hi-C data of two different cell lines (human frontal cortex and Drosophila male Dm-BG3c2 (BG3) cells), including a single-cell Hi-C dataset and the a corresponding bulk Hi-C dataset for each, were downloaded from the Gene Expression Omnibus (GEO) database (Barrett, et al., 2012; Edgar, et al., 2002). The single-cell Hi-C dataset and bulk Hi-C dataset of the human cell line (GEO no.: GSE130711) contains the data of 24 chromosomes (Chr. 1-22, X, and Y) (Lee, et al., 2019). The single-cell Hi-C dataset and corresponding bulk Hi-C dataset of Drosophila cell line (GEO no.: GSE131811) contain the data of 7 chromosomes (chr2L, chr2R, chr3L, chr3R, chr4, chrX, and chrM) (Ulianov, et al., 2021). Since both the human single-cell dataset and the Drosophila single-cell dataset contain the Hi-C data of many individual cells, the data of 3 randomly chosen individual cells from the human cell line and 2 randomly chosen individual cells from the Drosophila cell line were used to train and test the denoising methods.

The 40 kilobase (kb) resolution that can be well distinguished by the sparse single-cell Hi-C data was used to produce chromosomal contact matrices for both the single-cell Hi-C data and corresponding bulk Hi-C data. For the human cell line, the single-cell Hi-C data of one cell and the bulk Hi-C data for Chr. 1, 4, 6, 7, 9, 10, 13, 14, 15, 16, 17, 19, 20, and 22 are used as single-cell and bulk-cell training data respectively (called *human_cell_1_training_data* and *human_population_training_data*, respectively), those for Chr. 3, 5, 11, 21 are used as the single-cell and bulk-cell validation data (called *human_cell_1_validation_data and human_population_validation_data*, respectively), and those for Chr. 2, 8, 12, 18 are used as the single-cell and bulk-cell test dataset (called *human_cell_1_test_data* and *human_population_test_data*, respectively) to evaluate if the model can generalize from one chromosome to another one. Moreover, the single-cell data of another two randomly selected human cells are used as an additional single-cell test dataset (called *human_cells_2_3_test_data*) to check if the diffusion model can generalize from one cell to another one of the same species.

The single-cell Hi-C data of two randomly selected Drosophila cells and the corresponding bulk Hi-C data for 7 chromosomes (chr2L, chr2R, chr3L, chr3R, chr4, chrX, and chrM) are used as the single-cell and bulk-cell test datasets respectively (called *drosophila_cells_test_data* and *drosophia_population_test_data*, respectively) to evaluate if the diffusion model trained on the data of the human cell can generalize to Drosophila.

The original Hi-C chromosomal contact matrices in all the datasets downloaded above were normalized into the range [0, 1] first according to (Hong, et al., 2020) and then converted into the range [-1, 1] by the formula *y*_*out*_ = 2*y*_*input*_ − 1 because the diffusion process requires the input value is in the range [-1, 1] (Ho, et al., 2020). The normalized original data were treated as clean data (i.e., the ground truth labels). The normalized original chromosomal contact matrices were preprocessed by adding Gaussian noise (Dimmick, 2020; Kawar, et al., 2022) into them by the formula x = *y*_*out*_ + *Z* where 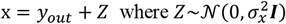 to generate noisy input data for denoising. *σ*_*x*_ is the noise level in the range [0, 1]. The higher the noise level in range [0, 1], the more noise is added, i.e., 0 means no Gaussian noise and 1 means full Gaussian noise. Two different noise levels (0.1 and 0.5) were used to generate noisy input data for the denoising tasks. For each noise level (0.1 or 0.5), there are 14 noisy chromosomal contact matrices in the single-cell or bulk training dataset (*human_cell_1_training_data* and *human_population_training_data*), 4 noisy chromosomal contact matrices in the single-cell or bulk validation dataset (*human_cell_1_validation_data and human_population_validation_data*), 4 noisy chromosomal contact matrices of Chr. 2, 8, 12, 18 in each of the two human single-cell test dataset (i.e., *human_cell_1_test_data* and *human_cells_2_3_test_data*), 7 noisy chromosomal contact matrices of chr2L, chr2R, chr3L, chr3R, chr4, chrX, and chrM in the Drosophila single-cell test dataset (i.e., *drosophia_cells_test_data*), 4 noisy chromosomal contact matrices of Chr. 2, 8, 12, 18 in the human bulk Hi-C test dataset (i.e., *human_population_test_data*), and 7 noisy chromosomal contact matrices of chr2L, chr2R, chr3L, chr3R, chr4, chrX, and chrM in Drosophila bulk Hi-C test dataset (*drosophila_population_test_data*).

64 × 64 sub-matrices were cropped from the noisy and clean chromosomal matrices in the training and validation datasets to create the inputs and labels to train and validate the methods. Training HiCDiff and DDPM was quite different from training the traditional non-diffusion deep learning methods. The non-diffusion methods were trained to predict clean data from noisy input data in the supervised mode. In contrast, the diffusion models (HiCDiff and DDPM) can be trained in both the unsupervised and unsupervised mode. In the unsupervised mode, they were fed with the clean data only, Gaussian noise was gradually added to the data in the forward diffusion process (**Algorithm 1**), and they were trained to recover the clean data iteratively in the reverse process (**Algorithm 2**). In the supervised mode, they were fed with both the clean data (labels) and down-sampled noisy data (i.e., condition) as input and were trained on them to learn to recover the clean data (**Algorithm 3**). During training, the validation datasets were used to inspect the convergence of the training process and select the best trained model for testing.

After the training, all the methods were blindly tested on the same test datasets to compare their performance. During the testing, a test matrix was divided into 64 × 64 sub-matrices with zero padding if necessary for a pretrained method to generate denoised submatrices. The denoised sub-matrices were then assembled into the full denoised matrix, which was compared with the ground truth (clean) matrix to evaluate its quality.

### 2.5 Evaluation Metrics

HiCDiff and DDPM diffusion models trained in the unsupervised mode (HiCDiff1 and DDPM1) and in the supervised mode (HiCDiff2 and DDPM2) were compared with several non-diffusion deep learning methods including DeepHiC (Hong, et al., 2020), HiCSR (Dimmick, 2020), SCHiCEDRN (Wang, et al., 2023), HiCPlus (Zhang, et al., 2018) and Loopenhance (DeepLoop) (Zhang, et al., 2022) in terms of multiple image-based evaluation metrics (PSNR: Peak signal-to-noise; SSIM: Structural Similarity Index Measure; MSE: Mean Squared Error; SNR: Signal-to-noise ratio), and the Hi-C reproducibility metric - the HiCRep score (Yang, et al., 2017) of measuring the similarity between denoised chromosomal contact matrices and clean matrices. To compare the denoised data generated by different methods against the clean data (target) after training, they were renormalized back into the same range [0, 1] according to (Kawar, et al., 2022) by the formula 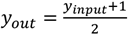, −1 ≤ *y*_*input*_ ≤ 1 before they were compared with the clean data that were also renormalized back into the range [0, 1] for the comparison because some image-based metrics such as SSIM are usually applied to non-negative matrices (Kawar, et al., 2022) and HiCRep (Yang, et al., 2017) requires two contact matrices in comparison to have non-negative values.

## 3. Results and Discussion

### 3.1 Diffusion models substantially improve the quality of single-cell Hi-C data

The quality of the chromosomal contact matrices denoised by the unsupervised and supervised diffusion models (HiCDiff1, DDPM1, HiCDiff2, and DDPM2) and that of the input matrices in the *human_cell_1_test_data* at the two noise levels are first compared in **Table 1**. All the four diffusion models substantially improve the quality of the input data according to the four image-based evaluation metrics. For instance, at the noise level of 0.1, the unsupervised HiCDiff1 increases PSNR from 28.9701 to 42.4171, SSIM from 0.1885 to 0.9662, and SNR from 27289 to 119604, and reduces MSE from 0.0013 to 0.00006; and at the higher noise level of 0.5, the improvement is generally more pronounced. Among the four diffusion models, the unsupervised diffusion models (HiCDiff1 and DDPM1) work better than their supervised counterparts (HiCDiff2 and DDPM2) respectively, indicating that the unsupervised (unconditional) mode is generally more effective than the supervised (conditional) mode for denoising the single-cell Hi-C data. Moreover, HiCDiff1 (or HiCDiff2) generally performs better than DDPM1 (or DDPM2), showing that the residual network architecture used by HiCDiff is more effective than the U-Net used by DDPM. Among the four diffusion models, the unsupervised HiCDiff (HiCDiff1) performs best on this dataset.

**Table 1.**
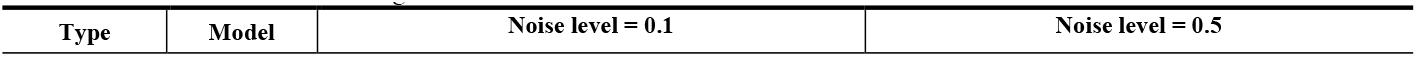
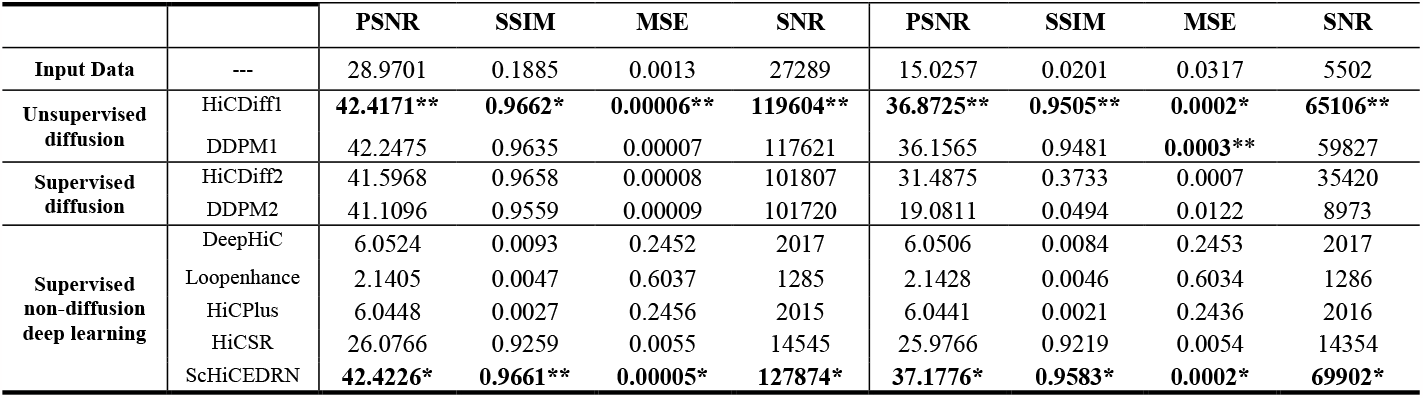
The results of the four unsupervised and supervised HiCDiff and DDPM diffusion models as well as five supervised non-diffusion deep learning methods on the *human_cell_1_test_data* at two input noise levels (0.1 and 0.5). The results of the input data without denoising are also shown as the baseline. “*” and “**” denote the best and second-best results, respectively. The unsupervised HiCDiff1 and DDPM1 used the linear noise variance schedule in its forward process, while the supervised HiCDiff2 and DDPM2 used the sigmoid noise variance schedule.

|  |  | PSNR | SSIM | MSE | SNR | PSNR | SSIM | MSE | SNR |
| --- | --- | --- | --- | --- | --- | --- | --- | --- | --- |
| **Input Data** | --- | 28.9701 | 0.1885 | 0.0013 | 27289 | 15.0257 | 0.0201 | 0.0317 | 5502 |
| **Unsupervised diffusion** | HiCDiff1 | **42.4171\*\*** | **0.9662\*** | **0.00006\*\*** | **119604\*\*** | **36.8725\*\*** | **0.9505\*\*** | **0.0002\*** | **65106\*\*** |
|  | DDPM1 | 42.2475 | 0.9635 | 0.00007 | 117621 | 36.1565 | 0.9481 | **0.0003\*\*** | 59827 |
| **Supervised diffusion** | HiCDiff2 | 41.5968 | 0.9658 | 0.00008 | 101807 | 31.4875 | 0.3733 | 0.0007 | 35420 |
|  | DDPM2 | 41.1096 | 0.9559 | 0.00009 | 101720 | 19.0811 | 0.0494 | 0.0122 | 8973 |
| **Supervised non-diffusion deep learning** | DeepHiC | 6.0524 | 0.0093 | 0.2452 | 2017 | 6.0506 | 0.0084 | 0.2453 | 2017 |
|  | Loopenhance | 2.1405 | 0.0047 | 0.6037 | 1285 | 2.1428 | 0.0046 | 0.6034 | 1286 |
|  | HiCPlus | 6.0448 | 0.0027 | 0.2456 | 2015 | 6.0441 | 0.0021 | 0.2436 | 2016 |
|  | HiCSR | 26.0766 | 0.9259 | 0.0055 | 14545 | 25.9766 | 0.9219 | 0.0054 | 14354 |
|  | ScHiCEDRN | **42.4226\*** | **0.9661\*\*** | **0.00005\*** | **127874\*** | **37.1776\*** | **0.9583\*** | **0.0002\*** | **69902\*** |

The similar results are observed on the single-cell Hi-C data of two human cells 2 and 3 in *human_cells_2_3_test* _*data* and the two Drosophila cells in *drosophia_cells_test_data* (supplemental **Table S1** and **S2**). Because the human cells 2 and 3 as well as the Drosophila cells were not used in the training at all, the results show that the diffusion models trained on one cell can work well on other cells of both the same species and different species and therefore are highly generalizable.

We also compare the four diffusion models trained on the bulk Hi-C data on the bulk Hi-C test datasets (*human_population_test data* and *drosophila_population_test_data*) (supplemental **Table S3** and **Table S4**). At the high noise level of 0.5, the unsupervised HiCDiff1 still performs best on both datasets. At the low noise level of 0.1, the supervised HiCDiff2 works best on *human_population_test_data*, while unsupervised HiCDiff1 works best on *drosophila_population_test_data*. Considering all the situations together, the unsupervised HiCDiff1 still performs best on the population Hi-C Data. It is worth noting that the amount of the improvement of the best diffusion model over the input data on the bulk Hi-C data (supplemental **Table S3** and **Table S4**) is generally lower than on the single-cell Hi-C data (supplemental **Table S1** and **Table S2**) probably because the single-cell Hi-C data is much sparser than the bulk Hi-C data and therefore has a larger room of improvement.

### 3.2 Unsupervised HiCDiff outperforms most supervised methods and is comparable to the best supervised method for denoising Hi-C data in terms of image-based metrics

The results of the diffusion models are also compared with the other five supervised non-diffusion deep learning methods (DeepHiC, hiCSR, HiCPlus, Loopenhance and ScHiCEDRN) on the single-cell Hi-C data of Human cell 1 (*human_cell_1_test_data*) in terms of the image-based evaluation metrics (**Table 1)** for two input noise levels: 0.1 and 0.5. The unsupervised HiCDiff (HiCDiff1) substantially outperforms all the supervised non-diffusion methods (e.g., DeepHiC, Loopenhance, HiCPlus and HiCSR) except ScHiCEDRN, while its performance is close to ScHiCEDRN. The similar results are observed on the two other human cells in *human_cells_2_3_test* (supplemental **Table S1)** as well as on the two Drosophila cells in *drosophia_cells_test_data* (supplemental **Table S2**). One unique feature of the unsupervised diffusion model like HiCDiff1 is that it only needs the clean data for training, which is different from the supervised methods that require pairs of clean and noisy data for training.

**Fig. 3** visually compares the chromosome matrices denoised by all the methods for the region (102.40-104.96 Mb) of Chr. 2 (**Fig. 3A**) and the region (307.20-309.76 Mb) of Chr. 12 (**Fig. 3B**). The sub-matrix with the highest contrast among these methods is highlighted by a green square in the first row of the matrices in **Fig. 3A** or **Fig. 3B**. The highlighted sub-matrix is enlarged in the second row in each sub figure, which shows that the data denoised by the diffusion models (HiCDiff and DDPM) are more similar to the target (clean) matrix than DeepHiC, hiCSR, HiCPlus and Loopenhance, while HiCDiff1 performs comparably to ScHiCEDRN.

**Fig. 3.**
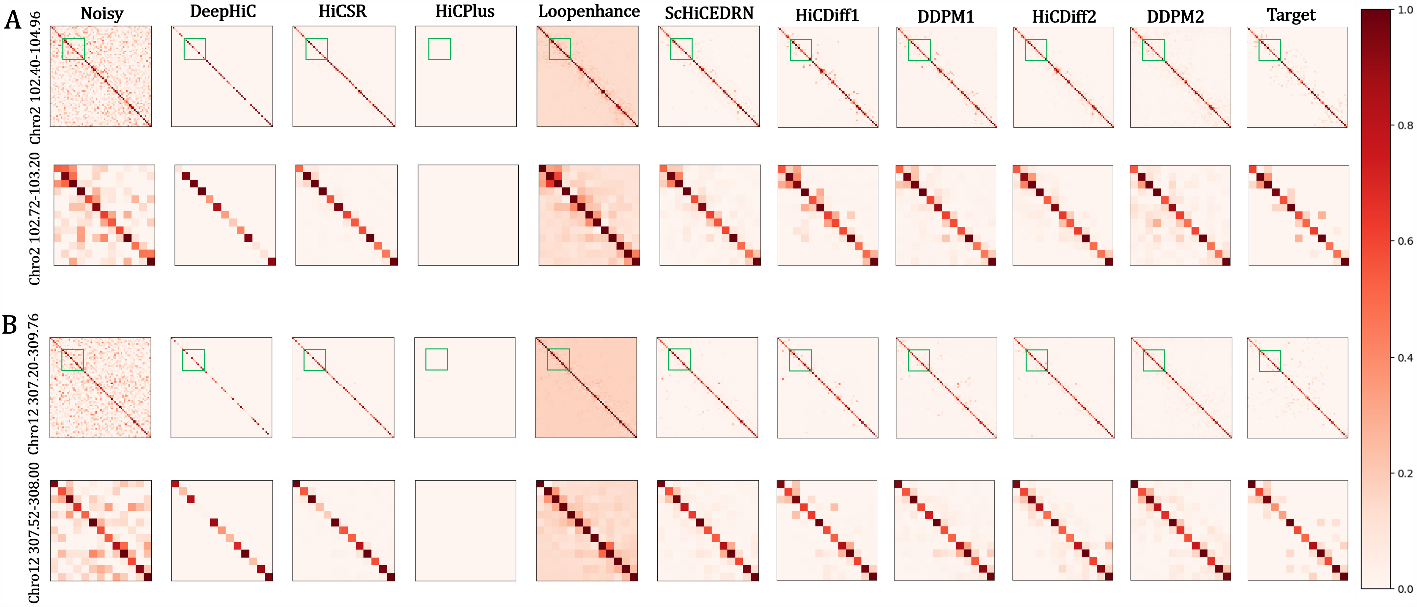
The heatmaps visualize the contact matrices denoised by the 9 denoising methods for Chr. 2 and Chr. 12 in *human_cell_1_test_data*. (**A**) The matrices of region 102.40 – 104.96 Mb of Chr. 2. The noisy input matrix and the clean target matrix are visualized at the beginning and the end, respectively. (**B**) The matrices for the region 307.20-309.76 Mb of Chr. 12. In the first row in (A) and (B), the green squares highlight the sub-region (102.72-103.20 Mb) of Chr. 2 and (307.52-308.00 Mb) of Chr. 12 with more pronounced difference between the matrices, which are enlarged in the second row.

In addition to comparing the diffusion methods with the supervised non-diffusion methods on the single-cell Hi-C data, we also trained them on the human bulk Hi-C training data and then tested them on the two bulk Hi-C test datasets (*human_population_test_data* and *drosophia_population_test_data*) (see results in **Table S3** and **S4)**. Largely similar results are seen on the bulk Hi-C data as on the single-cell Hi-C data. The unsupervised or supervised HiCDiff performs generally better than almost all the supervised non-diffusion methods (DeepHiC, HiCPlus, Loopenhance, and HiCSR) but a little worse than ScHiCEDRN. The results demonstrate that the diffusion models are also effective in denoising bulk Hi-C data.

### 3.3 HiCDiff achieves the state-of-the-art performance in terms of Hi-C reproducibility metric

HiCRep (Yang, et al., 2017) is a metric for systematically assessing the reproducibility of Hi-C data to capture the spatial features such as distance dependance or domain structure that are often neglected by other evaluation metrics. The higher HiCRep scores, the better the denoised Hi-C data are. In terms of the metric, we evaluated all the methods on the data of test chromosomes of the same human cell (*human_cell_1_test_data*) (**Fig. 4**), the data of the two unseen human cells in *human_cells_2_3_test_data* (**Fig. 5**), and two unseen Drosophila cells in *drosophia_cells_test_data* (**Fig. 5**).

**Fig. 4.**
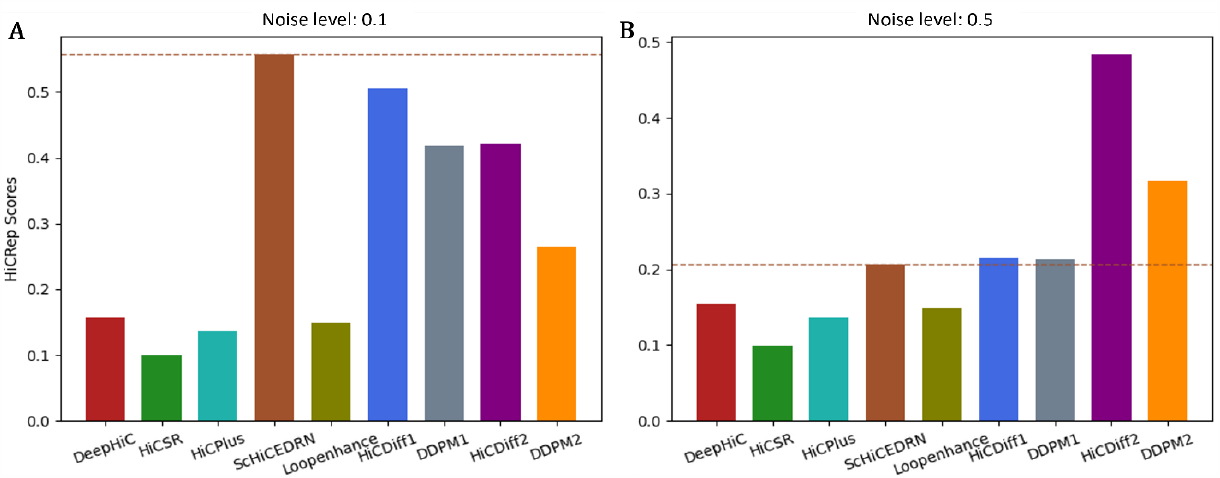
The box plot of the average HiCRep scores on *human_cell_1_test_data* at two different input noise levels (0.1 and 0.5). (**A**) Input noise level 0.1, (**B**) Input noise level 0.5.

**Fig. 5.**
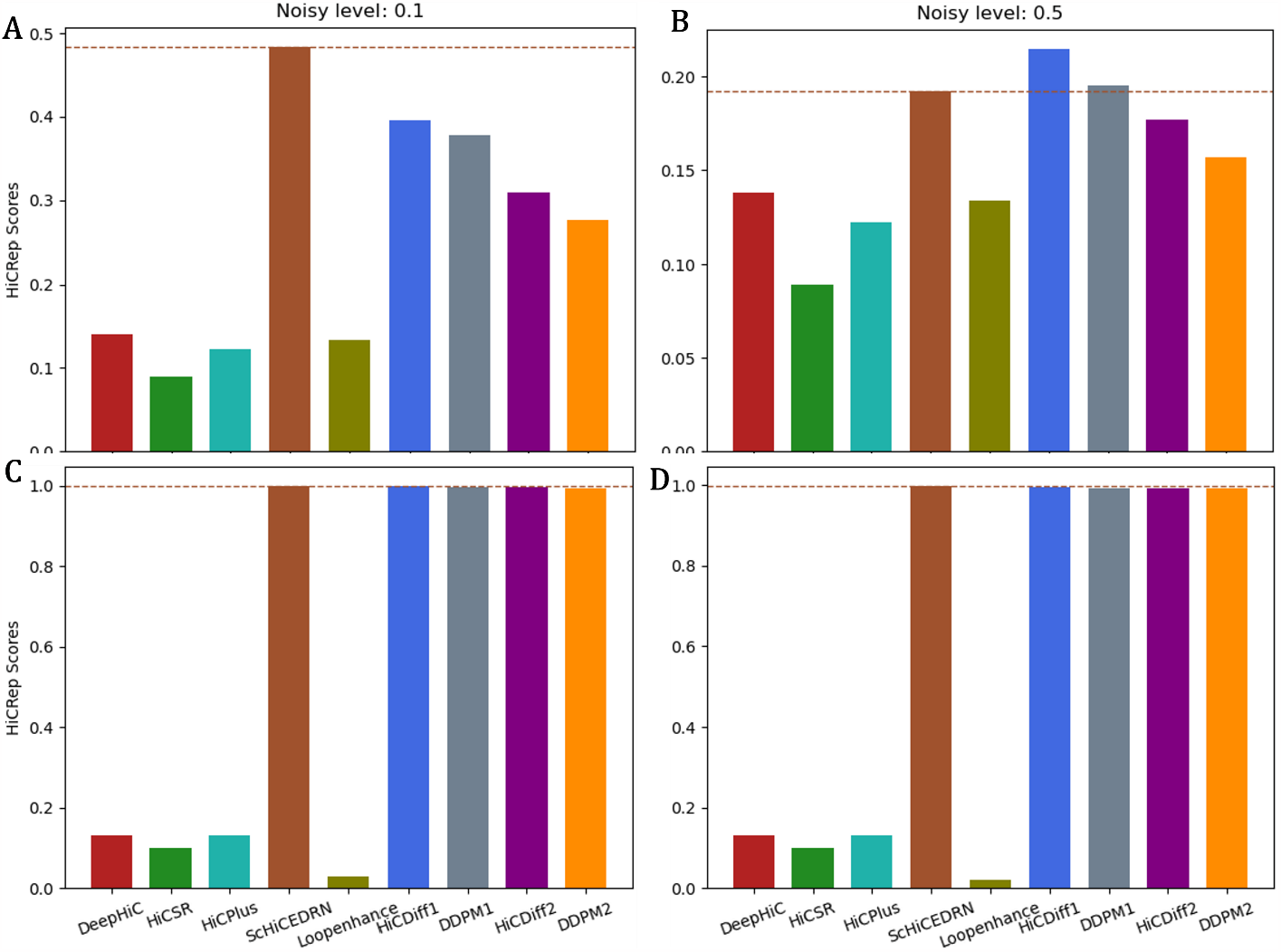
The box plot of the average HiCRep scores on human cells 2 and 3 of *human_cells_2_3_test_data* and Drosophila cells 1 and 2 of *drosophia_cells_test_data* at two noise levels. (**A**) On *human_cells_2_3_test_data* at input noise level 0.1, (**B**) On *human_cells_2_3_test_data* at input noise level 0.5. (**C**) On *drosophia_cells_test_data* at input noise level 0.1, and (**D**) On *drosophia_cells_test_data* at input noise level 0.5.

On the test data of the same human cell 1 (**Fig. 4A**) at the lower input noise level of 0.1, the performance of the unsupervised HiCDiff (HiDiff1) is second to a supervised method - ScHiCEDRN and much better than the other four supervised non-diffusion methods (DeepHiC, HiCPlus, Loopenhance, and HiCSR); at the higher input noise level of 0.5 (**Fig. 4B**), the supervised HiCDiff (HiCDiff2) performs much better than ScHiCEDRN and the other non-diffusion supervised methods.

On *human_cells_2_3_test_data*, at the lower input noise level 0.1 (**Fig. 5A**), ScHiCEDRN performs best, HiCDiff1 works the second best, and the diffusion methods (HiCDiff1, HiCDiff2, DDPM1, ad DDPM2) outperforms the other four supervised non-diffusion methods (DeepHiC, HiCPlus, Loopenhance, and HiCSR); at the higher input noise level 0.5 (**Fig. 5B**), the supervised HiCDiff2 performs best among all the methods.

On the test data of the *drosophia_cells_test_data*, at the two noise levels (0.1 and 0.5) (**Fig. 5C** and **Fig. 5D**), the four diffusion models (HiCDiff1, DDPM1, HiCDiff2, and DDPM2) and ScHiCEDRN have the similar performance, which is much better than that of the other four supervised non-diffusion methods.

We also evaluated all the methods on the human bulk Hi-C data in terms of HiCRep scores (**Fig. S1**). The four unsupervised/supervised diffusion models (HiCDiff1, DDPM1, HiCDidff1 and DDPM2) and two supervised deep learning methods (ScHiCEDRN and HiCSR) performs similarly at the two input noise levels, and they substantially outperform the other three non-diffusion methods (DeepHiC, HiCPlus, and Loopenhance).

Considering all the results above together, all the methods generally have higher HiCRep scores on the bulk Hi-C data than on the single-cell Hi-C data, which may be due to the high sparsity of the single-cell Hi-C data. On the single-cell Hi-C data, at higher input noise level (0.5), the supervised HiCDiff2 performs best and ScHiCEDRN second, while at lower input noise level (0.1), ScHiCEDRN has the best performance and the unsupervised HiCDiff1 the second.

### 3.4 The impact of the noise variance schedule on the performance of HiCDiff

The noise variance schedule (*β*_*t*_, 1 ≤ *t* ≤ *T*) in the diffusion model that controls how to gradually add Gaussian noise to the data in the forward diffusion process can influence its performance. We conducted an ablation study to investigate its impact on HiCDiff’s performance. The results of two different variance schedules (linear, sigmoid) on the *human_cell_1_test_data* at the input noise level of 0.1 are reported in **Table 2**. In the unsupervised mode, the linear variance schedule performs obviously better than the sigmoid variance schedule, while in the supervised mode, the sigmoid variance schedule performs slightly better than the linear variance schedule in terms of PSNR and SSIM, equally in terms of MSE, and slightly worse in terms of SNR. Overall, HiCDiff with the linear schedule in the unsupervised works best on the single-cell Hi-C dataset.

**Table 2.** The performance of the HiCDiff with the linear and sigmoid noise variance schedules on *human_cell_1_test_data* at the input noise level of 0.1. “*” and “**” denote the best and second-best results, respectively.

|  |  |  |  |  |  |
| --- | --- | --- | --- | --- | --- |
| Type | Model | PSNR | SSIM | MSE | SNR |
| Unsupervised, linear variance | HiCDiff1 | **42.4171\*** | **0.9662\*** | **0.000062\*** | **119604.93\*** |
| Supervised, linear variance | HiCDiff2 | 41.1033 | 0.9644 | **0.00008\*\*** | **104290.1\*\*** |
| Unsupervised, sigmoid variance | HiCDiff1 | 37.2066 | 0.7583 | 0.000191 | 70755.6 |
| Supervised, sigmoid variance | HiCDiff2 | **41.5968\*\*** | **0.9658\*\*** | **0.00008\*\*** | 101806.83 |

The results of the different variance schedule on the bulk Hi-C data (*human_population_test_data)* at the input noise level of 0.1 are reported in **Table S5**. Similarly, HiCDiff with the linear variance schedule performs better than sigmoid variance schedule in the unsupervised mode, while the situation is opposite in the supervised mode.

## 4. Conclusion

We developed the first diffusion models (HiCDiff and DDPM) to denoise single-cell Hi-C data in both unsupervised and supervised learning modes. The diffusion model can substantially improve the quality of the noisy Hi-C data. HiCDiff based on the residual network generally performs better than DDPM based on the standard U-Net in terms of multiple metrics. On multiple test datasets of two input noise levels from the same cell, different single cells, cells of different species, the unsupervised HiCDiff achieved the performance similar to the state-of-the-art supervised ScHiCEDRN and better than all other non-diffusion supervised methods, demonstrating that diffusion models are a useful approach to the single-cell Hi-C data denoising problem. Its performance generalizes well from one cell to another cell and from one species to another species. Moreover, HiCDiff performs well in denoising bulk Hi-C data.

## Code availability

The source code of HiCDiff is available at the GitHub repository: https://github.com/BioinfoMachineLearning/hicdiff.

## Data availability

The source code of ScHiCEDRN is available at the GitHub repository: https://github.com/BioinfoMachineLearning/hicdiff. The pyTorch version of other four methods DeepHiC, HiCPlus, HiCSR and Loopenhance can be accessed at https://github.com/omegahh/DeepHiC, https://github.com/wangjuan001/hicplus, https://github.com/PSI-Lab/HiCSR, and https://github.com/JinLabBioinfo/DeepLoop, respectively.

The original training, validation and test datasets in our experiments are Cooler files. The Cooler file containing both the single-cell Hi-C data of the three human cells (GEO accession number: GSE130711) and population human Hi-C data (GEO accession number: GSE130711) were downloaded from https://salkinstitute.app.box.com/s/fp63a4j36m5k255dhje3zcj5kfuzkyj1. The Cooler file containing both the single-cell Hi-C data of two Drosophila cells (GEO accession number: GSE131811) and population Drosophila Hi-C data (GEO accession number: GSE131811) were obtained from https://www.ncbi.nlm.nih.gov/geo/query/acc.cgi?acc=GSE131811. All the original and processed datasets can also be downloaded from Zenodo: https://doi.org/10.5281/zenodo.10223407.

## Authors’ Contributions

JC conceived the project. YW and JC designed the methods. YW implemented the methods, performed the experiments, and collected and analyzed the data. YW and JC wrote the manuscript.

## Acknowledgements

This work was partially supported by the William and Nancy Thompson Professorship and a Curators’ Professorship to JC.

## Notes

### Competing Interest Statement

The authors have declared no competing interest.

## References

Bardsley, J.M. MCMC-based image reconstruction with uncertainty quantification. SIAM Journal on Scientific Computing 2012;34(3):A1316–A1332.

Barrett, T., et al. NCBI GEO: archive for functional genomics data sets—update. Nucleic acids research 2012;41(D1):D991–D995.

Dimmick, M. HiCSR: a Hi-C super-resolution framework for producing highly realistic contact maps. University of Toronto (Canada); 2020.

Dixon, J.R., et al. Topological domains in mammalian genomes identified by analysis of chromatin interactions. Nature 2012;485(7398):376–380.

Edgar, R., Domrachev, M. and Lash, A.E. Gene Expression Omnibus: NCBI gene expression and hybridization array data repository. Nucleic acids research 2002;30(1):207–210.

Goodfellow, I., et al. Generative adversarial nets. Advances in neural information processing systems 2014;27.

Guo, Z., et al. Diffusion models in bioinformatics and computational biology. Nature Reviews Bioengineering 2023:1–19.

Hicks, P. and Oluwadare, O. HiCARN: resolution enhancement of Hi-C data using cascading residual networks. Bioinformatics 2022;38(9):2414–2421.

Highsmith, M. and Cheng, J. Vehicle: a variationally encoded hi-c loss enhancement algorithm for improving and generating hi-c data. Scientific Reports 2021;11(1):8880.

Ho, J., Jain, A. and Abbeel, P. Denoising diffusion probabilistic models. Advances in Neural Information Processing Systems 2020;33:6840–6851.

Hong, H., et al. DeepHiC: A generative adversarial network for enhancing Hi-C data resolution. PLoS computational biology 2020;16(2):e1007287.

Karras, T., et al. Progressive growing of gans for improved quality, stability, and variation. arXiv preprint arXiv:1710.10196 2017.

Kawar, B., et al. Denoising diffusion restoration models. arXiv preprint arXiv:2201.11793 2022.

Lee, D.-S., et al. Simultaneous profiling of 3D genome structure and DNA methylation in single human cells. Nature methods 2019;16(10):999–1006.

Lehtinen, J., et al. Noise2Noise: Learning image restoration without clean data. arXiv preprint arXiv:1803.04189 2018.

Li, W., et al. Hi-Corrector: a fast, scalable and memory-efficient package for normalizing large-scale Hi-C data. Bioinformatics 2015;31(6):960–962.

Lieberman-Aiden, E., et al. Comprehensive mapping of long-range interactions reveals folding principles of the human genome. science 2009;326(5950):289–293.

Nagano, T., et al. Single-cell Hi-C reveals cell-to-cell variability in chromosome structure. Nature 2013;502(7469):59–64.

Nagano, T., et al. Cell-cycle dynamics of chromosomal organization at single-cell resolution. Nature 2017;547(7661):61–67.

Nora, E.P., et al. Spatial partitioning of the regulatory landscape of the X-inactivation centre. Nature 2012;485(7398):381–385.

Ongie, G., et al. Deep learning techniques for inverse problems in imaging. IEEE Journal on Selected Areas in Information Theory 2020;1(1):39–56.

Pal, K., Forcato, M. and Ferrari, F. Hi-C analysis: from data generation to integration. Biophysical reviews 2019;11:67–78.

Saharia, C., et al. Image super-resolution via iterative refinement. IEEE Transactions on Pattern Analysis and Machine Intelligence 2022.

Servant, N., et al. HiC-Pro: an optimized and flexible pipeline for Hi-C data processing. Genome biology 2015;16(1):1–11.

Sohl-Dickstein, J., et al. Deep unsupervised learning using nonequilibrium thermodynamics. In, International Conference on Machine Learning. PMLR; 2015. p. 2256–2265.

Tan, L., et al. Three-dimensional genome structures of single diploid human cells. Science 2018;361(6405):924–928.

Trieu, T. and Cheng, J. MOGEN: a tool for reconstructing 3D models of genomes from chromosomal conformation capturing data. Bioinformatics 2016;32(9):1286–1292.

Ulianov, S.V., et al. Order and stochasticity in the folding of individual Drosophila genomes. Nature communications 2021;12(1):41.

Wang, Y., Guo, Z. and Cheng, J. Single-cell Hi-C data enhancement with deep residual and generative adversarial networks. Bioinformatics 2023;39(8):btad458.

Wolff, J., Backofen, R. and Grüning, B. Loop detection using Hi-C data with HiCExplorer. Gigascience 2022;11:giac061.

Yang, T., et al. HiCRep: assessing the reproducibility of Hi-C data using a stratum-adjusted correlation coefficient. Genome research 2017;27(11):1939–1949.

Zhang, K., et al. Beyond a gaussian denoiser: Residual learning of deep cnn for image denoising. IEEE transactions on image processing 2017;26(7):3142–3155.

Zhang, R., Zhou, T. and Ma, J. Multiscale and integrative single-cell Hi-C analysis with Higashi. Nature biotechnology 2022;40(2):254–261.

Zhang, S., et al. DeepLoop robustly maps chromatin interactions from sparse allele-resolved or singlecell Hi-C data at kilobase resolution. Nature Genetics 2022;54(7):1013–1025.

Zhang, Y., et al. Enhancing Hi-C data resolution with deep convolutional neural network HiCPlus. Nature communications 2018;9(1):750.

