## supplemental information for "HiCDiff: single-cell Hi-C data denoising with diffusion models"

| **Table S1**. The results of the four unsupervised and supervised HiCDiff and DDPM diffusion models as well as five other non-diffusion supervised deep learning methods for denoising one single-cell Hi-C data of *human cells 2 and 3 on human_cells_2_3_test_data* at two input noise levels (0.1 and 0.5). The results of the input data without denoising are also shown as the baseline. “*” and “**” denote the best and second-best results, respectively. The unsupervised HiCDiff1 and DDPM1 used the linear noise variance schedule in its forward process, while the supervised HiCDiff2 and DDPM2 used the sigmoid noise variance schedule. | | | | | | | | | | | | | | | |
| --- | --- | --- | --- | --- | --- | --- | --- | --- | --- | --- | --- | --- | --- | --- | --- |
| **Type** | **Model** | **Noise level = 0.1** | | | | | | **Noise level = 0.5** | | | | | | | |
|  |  | **PSNR** | **SSIM** | **MSE** | | **SNR** | | **PSNR** | | **SSIM** | | **MSE** | | | **SNR** |
| **Input Data** | --- | 28.9786 | 0.1868 | 0.0013 | | 30275 | | 15.0271 | | 0.0193 | | 0.0324 | | | 6097 |
| **Unsupervised diffusion** | HiCDiff1 | **42.9328**** | **0.9699*** | **0.00005*** | | **144480**** | | **36.8585**** | | **0.9591**** | | **0.000225**** | | | **75193 **** |
|  | DDPM1 | 42.7655 | 0.9680 | **0.00006**** | | 138340 | | 36.1697 | | 0.9533 | | 0.0003 | | | 67284 |
| **Supervised diffusion** | HiCDiff2 | 42.0023 | 0.9698 | 0.000075 | | 118891 | | 35.4670 | | 0.9585 | | 0.00031 | | | 61838 |
|  | DDPM2 | 41.5441 | 0.9598 | 0.000085 | | 112110 | | 35.3751 | | 0.9523 | | 0.00039 | | | 56137 |
| **Supervised non-diffusion deep learning** | DeepHiC | 6.0564 | 0.0100 | 0.2391 | | 1992 | | 6.0547 | | 0.0092 | | 0.2392 | | | 2396 |
|  | Loopenhance | 2.1454 | 0.0059 | 0.5885 | | 1582 | | 2.1472 | | 0.0058 | | 0.5883 | | | 1528 |
|  | HiCPlus | 6.0483 | 0.0036 | 0.2438 | | 2903 | | 6.0398 | | 0.0035 | | 0.2514 | | | 2915 |
|  | HiCSR | 25.9040 | 0.9298 | 0.0075 | | 16360 | | 25.8143 | | 0.9189 | | 0.0078 | | | 16289 |
|  | ScHiCEDRN | **43.2613*** | **0.9697**** | **0.00005*** | | **147489*** | | **37.4017*** | | **0.9624*** | | **0.00019*** | | | **78468*** |
| **Table S2**. The results of the four unsupervised and supervised HiCDiff and DDPM diffusion models as well as five other supervised non-diffusion deep learning methods for denoising one individual single-cell Hi-C data of *drosophila cells 1 and 2 on drosophia_cells_test_data* at two input noise levels (0.1 and 0.5). The results of the input data without denoising are also shown as the baseline. “*” and “**” denote the best and second-best results, respectively. The unsupervised HiCDiff1 and DDPM1 used the linear noise variance schedule in its forward process, while the supervised HiCDiff2 and DDPM2 used the sigmoid noise variance schedule. | | | | | | | | | | | | | | | |
| **Type** | **Model** | **Noise level = 0.1** | | | | | | | **Noise level = 0.5** | | | | | | |
|  |  | **PSNR** | **SSIM** | | **MSE** | | **SNR** | | **PSNR** | | **SSIM** | | **MSE** | **SNR** | |
| **Input Data** | --- | 28.4453 | 0.2325 | | 0.0015 | | 91034 | | 14.5864 | | 0.0356 | | 0.0354 | 18482 | |
| **Unsupervised diffusion** | HiCDiff1 | **41.3910**** | **0.9729*** | | **0.00007*** | | **427813**** | | **34.2649*** | | **0.9482**** | | **0.0004*** | **180594**** | |
|  | DDPM1 | 41.0571 | 0.9606 | | 0.00018 | | 402197 | | 31.9191 | | 0.9343 | | 0.0007 | 137326 | |
| **Supervised diffusion** | HiCDiff2 | 38.9784 | **0.9656**** | | **0.00016**** | | 311240 | | 32.1794 | | 0.9426 | | **0.0006**** | 144410 | |
|  | DDPM2 | 37.2031 | 0.9554 | | 0.00025 | | 249838 | | 31.2399 | | 0.9344 | | 0.0010 | 125422 | |
| **Supervised non-diffusion deep learning** | DeepHiC | 6.6774 | 0.1249 | | 0.2092 | | 7517 | | 6.6766 | | 0.1238 | | 0.2092 | 7516 | |
|  | Loopenhance | 2.7553 | 0.1147 | | 0.5162 | | 4786 | | 2.7626 | | 0.1127 | | 0.3523 | 4790 | |
|  | HiCPlus | 6.6765 | 0.1231 | | 0.2093 | | 5554 | | 6.6494 | | 0.1218 | | 0.2043 | 7089 | |
|  | HiCSR | 14.9658 | 0.7941 | | 0.0397 | | 19306 | | 14.4831 | | 0.7793 | | 0.0448 | 18640 | |
|  | ScHiCEDRN | **41.6869*** | **0.9729*** | | **0.00007*** | | **433829*** | | **34.0881**** | | **0.9495*** | | **0.0004*** | **181713*** | |

| **Table S3**. The results of the four unsupervised and supervised HiCDiff and DDPM diffusion models as well as five other supervised non-diffusion deep learning methods for denoising population Hi-C data of *human_population_test_data data* at two input noise levels (0.1 and 0.5). The results of the input data without denoising are also shown as the baseline. “*” and “**” denote the best and second-best results, respectively. The unsupervised HiCDiff1 and DDPM1 used the linear noise variance schedule in its forward process, while the supervised HiCDiff2 and DDPM2 used the sigmoid noise variance schedule. | | | | | | | | | |
| --- | --- | --- | --- | --- | --- | --- | --- | --- | --- |
| **Type** | **Model** | **Noise level = 0.1** | | | | **Noise level = 0.5** | | | |
|  |  | **PSNR** | **SSIM** | **MSE** | **SNR** | **PSNR** | **SSIM** | **MSE** | **SNR** |
| **Input Data** | --- | 28.2283 | 0.8332 | 0.0015 | 1448099 | 14.5308 | 0.4198 | 0.0355 | 299486 |
| **Unsupervised diffusion** | HiCDiff1 | 27.6019 | 0.8967 | 0.0017 | 1348932 | **16.7427**** | **0.5031**** | **0.0215**** | **385109**** |
|  | DDPM1 | 26.1168 | 0.8804 | 0.0024 | 1131995 | 16.5844 | 0.4729 | 0.0223 | 378497 |
| **Supervised diffusion** | HiCDiff2 | **31.5966**** | **0.9766*** | **0.0008*** | **1911097*** | 16.5747 | 0.4954 | 0.0249 | 377742 |
|  | DDPM2 | 29.0613 | 0.9557 | 0.0012 | 1597274 | 15.3508 | 0.4384 | 0.0295 | 328521 |
| **Supervised non-diffusion deep learning** | DeepHiC | 7.7732 | 0.2839 | 0.1641 | 139409 | 7.6184 | 0.2070 | 0.1703 | 136823 |
|  | Loopenhance | 4.7532 | 0.2301 | 0.3299 | 98292 | 4.6358 | 0.1686 | 0.3391 | 96949 |
|  | HiCPlus | 7.7692 | 0.2744 | 0.1643 | 139339 | 6.8179 | 0.0659 | 0.2056 | 124487 |
|  | HiCSR | 30.8713 | 0.9697 | 0.0008* | 1901590 | 15.8383 | 0.4545 | 0.0263 | 347969 |
|  | ScHiCEDRN | **31.9009*** | **0.9748**** | **0.0005*** | **2038916*** | **16.9232*** | **0.5041*** | **0.0205*** | **393977*** |

| **Table S4**. The results of the four unsupervised and supervised HiCDiff and DDPM diffusion models as well as five other supervised non-diffusion deep learning methods for denoising population Hi-C data of *drosophia_population_test_data* at two input noise levels (0.1 and 0.5). The results of the input data without denoising are also shown as the baseline. “*” and “**” denote the best and second-best results, respectively. The unsupervised HiCDiff1 and DDPM1 used the linear noise variance schedule in its forward process, while the supervised HiCDiff2 and DDPM2 used the sigmoid noise variance schedule. | | | | | | | | | |
| --- | --- | --- | --- | --- | --- | --- | --- | --- | --- |
| **Type** | **Model** | **Noise level = 0.1** | | | | **Noise level = 0.5** | | | |
|  |  | **PSNR** | **SSIM** | **MSE** | **SNR** | **PSNR** | **SSIM** | **MSE** | **SNR** |
| **Input Data** | --- | 27.8107 | 0.5470 | 0.0017 | 245307 | 14.6267 | 0.0984 | 0.0348 | 54062 |
| **Unsupervised diffusion** | HiCDiff1 | **31.3634**** | **0.7598**** | **0.0008**** | **374926**** | **26.2581**** | **0.5909**** | **0.0033**** | **198208**** |
|  | DDPM1 | 30.3938 | 0.7058 | 0.0009 | 351776 | 25.2610 | 0.4070 | 0.0064 | 155862 |
| **Supervised diffusion** | HiCDiff2 | 28.9845 | 0.6319 | 0.0013 | 273396 | 23.3784 | 0.4617 | 0.0049 | 163345 |
|  | DDPM2 | 28.7563 | 0.6236 | 0.0014 | 266188 | 22.9597 | 0.4228 | 0.0055 | 156177 |
| **Supervised non-diffusion deep learning** | DeepHiC | 7.0799 | 0.1501 | 0.1915 | 23163 | 7.0685 | 0.1443 | 0.1921 | 23130 |
|  | Loopenhance | 4.2113 | 0.1171 | 0.3751 | 16530 | 4.3095 | 0.1017 | 0.3662 | 16731 |
|  | HiCPlus | 7.0788 | 0.1339 | 0.1916 | 23160 | 6.9511 | 0.1029 | 0.1977 | 22790 |
|  | HiCSR | 29.6669 | 0.4933 | 0.0012 | 294032 | 22.8838 | 0.3708 | 0.0054 | 137215 |
|  | ScHiCEDRN | **31.9009*** | **0.7689*** | **0.0007*** | **380340*** | **26.7303*** | **0.6110*** | **0.00239*** | **207566*** |

| **Table S5**. The impact of the variance schedule on the performance of HiCDiff on *human_population_test_data* at the input noise level of 0.1. | | | | | |
| --- | --- | --- | --- | --- | --- |
| **Type** | **Model** | **PSNR** | **SSIM** | **MSE** | **SNR** |
| Unsupervised, linear variance | HiCDiff1 | 27.6019 | 0.8967 | 0.0017 | 1348932.1 |
| Supervised, linear variance | HiCDiff2 | 29.6478 | 0.9625 | 0.0011 | 1708533.1 |
| Unsupervised, sigmoid variance | HiCDiff1 | 27.3687 | 0.9219 | 0.0018 | 1309345.1 |
| Supervised, sigmoid variance | HiCDiff2 | 31.5966 | 0.9766 | 0.0008 | 1911097.8 |

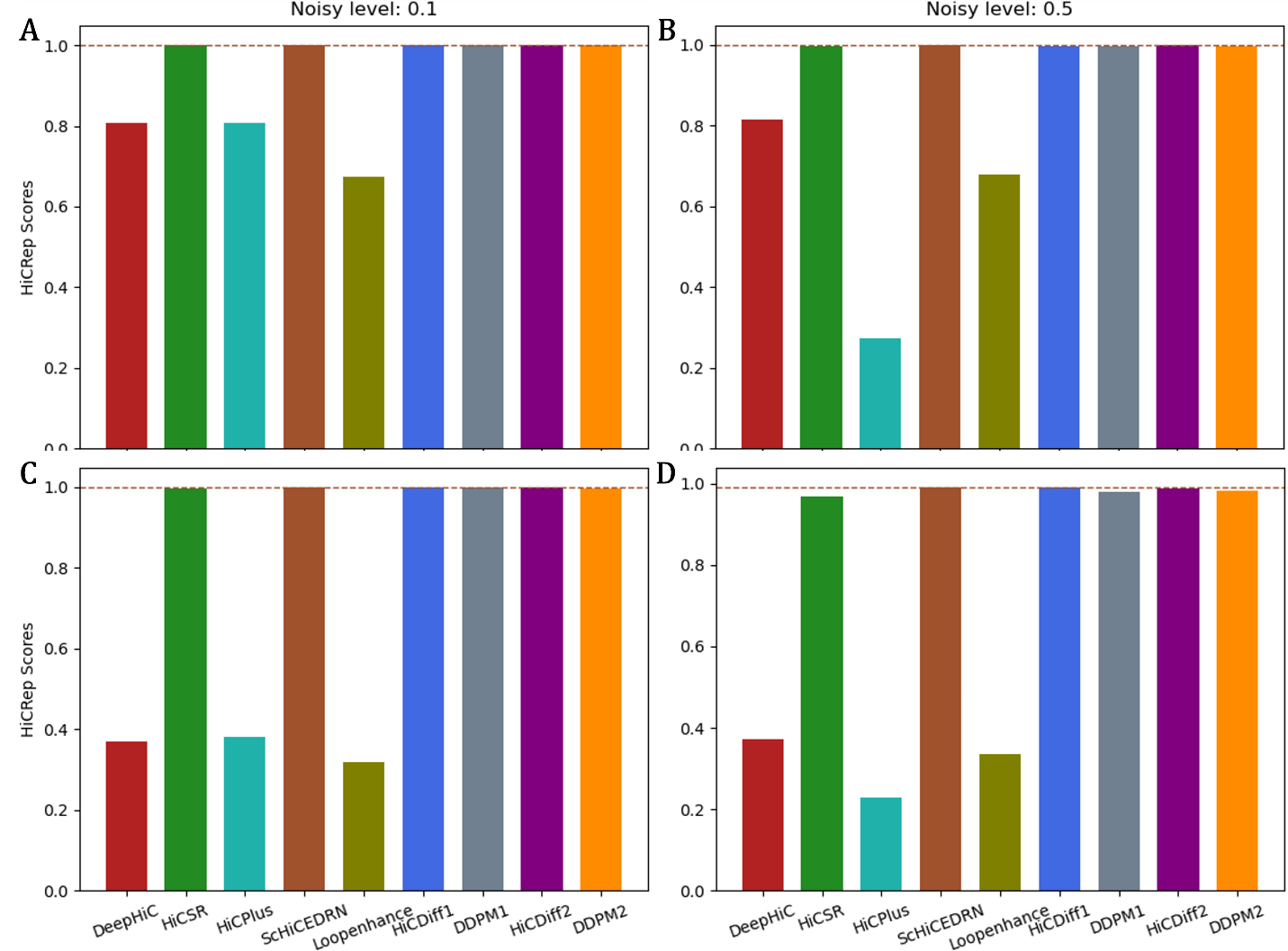

**Fig. S1.** The box plot of the average HiCRep scores from the *human_population_test_data* and *drosophila_population_test_data* Hi-C data at the different noise levels (0.1 and 0.5). (**A**) On human population cells at input noise level 0.1, (**B**) On human population cells at input noise level 0.5. (**C**) On Drosophila population cells at input noise level 0.1, and (**D**) On Drosophila population cells at input noise level 0.5.
